# Reconstructing temporal and spatial dynamics in single-cell experiments

**DOI:** 10.1101/697151

**Authors:** Karsten Kuritz, Daniela Stöhr, Daniela Maichl, Nadine Pollak, Markus Rehm, Frank Allgöwer

## Abstract

Modern cytometry methods allow collecting complex, multi-dimensional data sets from heterogeneous cell populations at single-cell resolution. While methods exist to describe the progression and order of cellular processes from snapshots of such populations, these descriptions are limited to arbitrary pseudotime scales. Here we describe MAPiT, an universal transformation method that recovers real-time dynamics of cellular processes from pseudotime scales. As use cases, we applied MAPiT to two prominent problems in the flow-cytometric analysis of heterogeneous cell populations: (1) recovering the kinetics of cell cycle progression in unsynchronized and thus unperturbed cell populations, and (2) recovering the spatial arrangement of cells within multi-cellular spheroids prior to spheroid dissociation for cytometric analysis. Since MAPiT provides a theoretic basis for the relation of pseudotime values to real temporal and spatial scales, it can be used broadly in the analysis of cellular processes with snapshot data from heterogeneous cell populations.

## Introduction

Quantitative single-cell measurements with ten to several thousands of cellular components in large populations provide new opportunities and challenges to study biological processes (Wagner et al., 2016; Regev et al., 2017; Pijuan-Sala et al., 2018). Since cells within heterogeneous populations span all stages and transitions of a biological process of interest and hence also the dynamics of cellular components, the temporal evolution of cellular signaling events can be inferred from static snapshot data. Several algorithms, such as Wanderlust, Monocle or diffusion pseudotime (DPT), designed to reconstruct cell trajectories, order single-cell data in *pseudotime* – a quantitative measure of progress through a biological process (Haghverdi, Buettner, et al., 2014; Bendall et al., 2014; Angerer et al., 2016; Trapnell et al., 2014; Qiu et al., 2017; Weinreb et al., 2018). However, these pseudotime trajectories may deviate substantially from the real-time trajectory (Yuan et al., 2017; Tanay and Regev, 2017). Alternative approaches attempting to transfer pseudotime to real-time analysis are technically restricted, e.g. limited to cell cycle analysis (Kafri et al., 2013; Kuritz et al., 2017), require specific source data, such as single-cell RNA-seq data (La Manno et al., 2018), or require computationally expensive estimations of non-indentifiable functions (Weinreb et al., 2018; Fischer et al., 2019). A more detailed discussion on attempts to transform pseudotime scales is provided in the Supplementary Information.

Here, we developed a measure-preserving transformation of pseudotime into real-time, a <u>MAP</u> of pseudotime into <u>T</u>ime, in short MAPiT. MAPiT generalizes approaches based on ergodic principles to provide a simple and at the same time universal method to obtain true scale dynamics from pseudotime ordering (Fig. 1).

**Figure 1:**
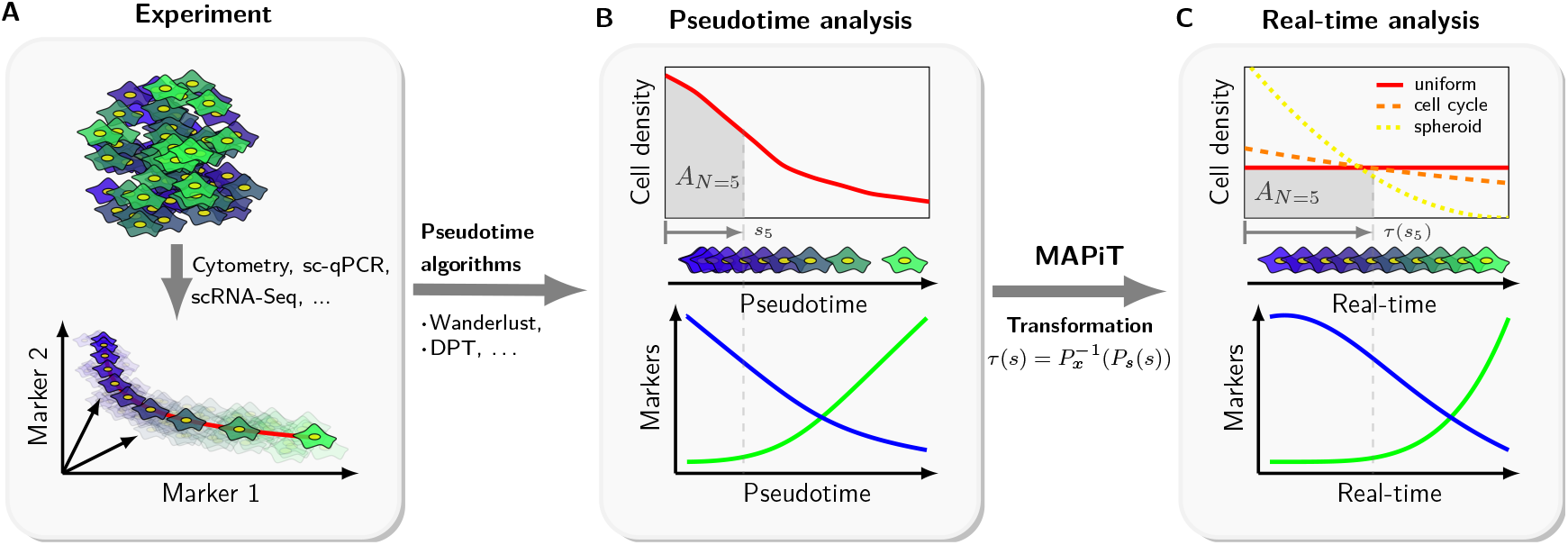
MAPiT deduces process dynamics from single-cell snapshot data. (**A**) Cells from single-cell experiments of a heterogeneous population are ordered on a process manifold in dataspace by pseudotime algorithms. (**B**) Cell density and marker trajectories on pseudotime scale vary with the distance measure used by the pseudotime algorithm and real temporal trajectories cannot be deduced. Cell density, order and trajectories for two markers on pseudotime scale are shown for an exemplary process. As an example pseudotime position of the fifth displayed cell *s*_5_ and associated area under the cell density curve *A*_*N*=5_ are indicated in gray. (**C**) Nonlinear transformation of pseudotime scale recovers true scale dynamics. MAPiT uses prior knowledge of cell densities on the real scale to transform pseudotime to real time by enforcing equality for the area under the density curves at corresponding points on both scales (gray areas). Cell order and marker trajectories are shown for an examplary uniform distribution on the real scale. Positions of cells across the cell cycle (dashed, orange) or decreasing number of cells towards the center of spheroid cultures (dotted, yellow) are other real scale densities.

## Results

Common pseudotime algorithms order cells on a pseudotime scale based on a distance metric in the data space, and this metric differs between algorithms (Saelens et al., 2019). Pseudotime values furthermore strongly depend on the measured cellular components. MAPiT resolves the arbitrariness of pseudotime by nonlinearly transforming pseudotime to the true scale of the process. This is based on a “measure-preserving transformation” which ensures that the area under the curve is conserved when transforming a probability distribution (see methods).

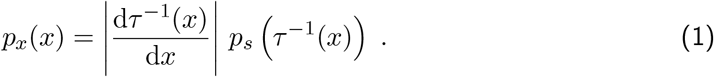

The mapping *τ*: *s* → *x* from pseudotime to real-time is obtained by solving equation (1) for *τ*, which then depends on the cumulative distributions of cells on both scales:

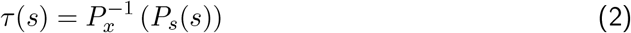

The transformation requires knowledge of the distributions (or cumulative distributions) of cells on both scales (pseudotime and desired scale). Pseudotime values from experimental data can be used to calculate the distribution on the pseudotime scale. In contrast, a priori knowledge of the process of interest must be used to derive the distribution of cells on the desired real-time scale, as we demonstrate in the following examples.

We first applied MAPiT to analyse cell cycle progression. To this end, we used a static flow cytometric measurement of DNA and *mAG-hGeminin (1-110)*, a fluorescent ubiquitination-based cell cycle indicator (Fig. 2 A) (Sakaue-Sawano et al., 2008; Kuritz et al., 2017), in unperturbed NCI-H460/geminin cells to reconstruct the kinetics of geminin. The pseudotime obtained with the markers was mapped to the temporal scale on which the the cell cycle progresses, namely the age of a cell (time since cytokinesis). To achieve this, MAPiT derives real-time trajectories from the steady state age distribution (Powell, 1956; Kafri et al., 2013; Kuritz et al., 2017) (Fig. 2 B). Temporal trajectories of geminin obtained with MAPiT corresponded excellently to geminin kinetics obtained by single cell time-lapse microscopy (Fig. 2 D), as examplified by the boost in geminin intensity at approximately 7 *h*, indicating the onset of S-phase. This result therefore highlights the temporal accuracy of the real-time scale obtained by MAPiT for cell cycle analysis based on snapshot flow cytometric data.

**Figure 2:**
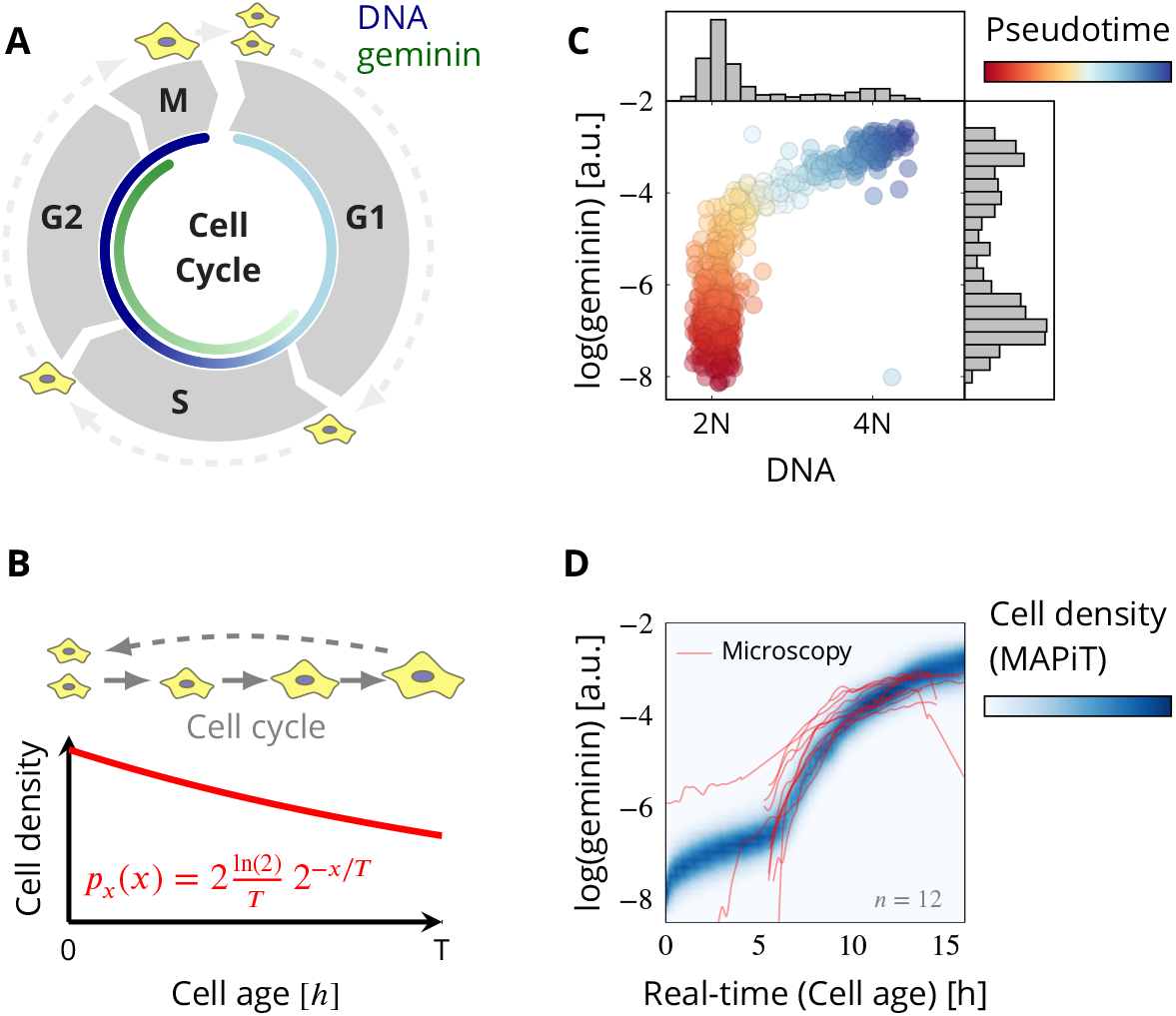
MAPiT recovers cell cycle dynamics. (**A**) Schematic of the cell cycle with geminin expression, a marker for cell cycle progression, starting at the onset of S phase. (**B**) MAPiT employs steady state age distribution of unsynchronized cell populations with cell cycle length *T*. (**C**) DNA and geminin signals from an unsynchronized population of NCI-H460/geminin cells were used to obtain a pseudotemporal ordering of the population. (**D**) Reconstructed temporal profile of geminin signal density obtained with MAPiT and single-cell trajectories from time-lapse imaging correlate strongly.

Multicellular spheroids grown from cancer cells are widely used as avascular tumor models (Vörsmann et al., 2013; Jabs et al., 2017; Lin and Chang, 2008). As a consequence of nutrient and oxygen deprivation within the spheroids, proliferative cells begin to enclose inner layers of quiescent and necrotic cells, resembling a zonation found in solid tumours (Freyer and Sutherland, 1986; LaBarbera et al., 2012). Current routine methods to study spatial distributions and patterns of cellular markers are restricted to intact spheroids, technically cumbersome and of limited throughput, since they rely on sequential spheroid fixation, sectioning and imaging procedures. By dissociating tumor spheroids for single cell experiments, spatial information across which cell-to-cell heterogeneities in tumor cells spheroids manifest is lost. We applied MAPiT to study if we can recover spatial scales from flow cytometric measurements of dissociated spheroids in a reliable and robust manner. For our studies, we grew spheroids of HCT116 cells to diameters of approximately 500 *μm*. Cells were stained for DNA as measure for cell cycle stage, RNA as indicator for transcriptional activity, Ki-67 as marker for proliferation and p27 as marker for quiescence (Fig. 3).

**Figure 3:**
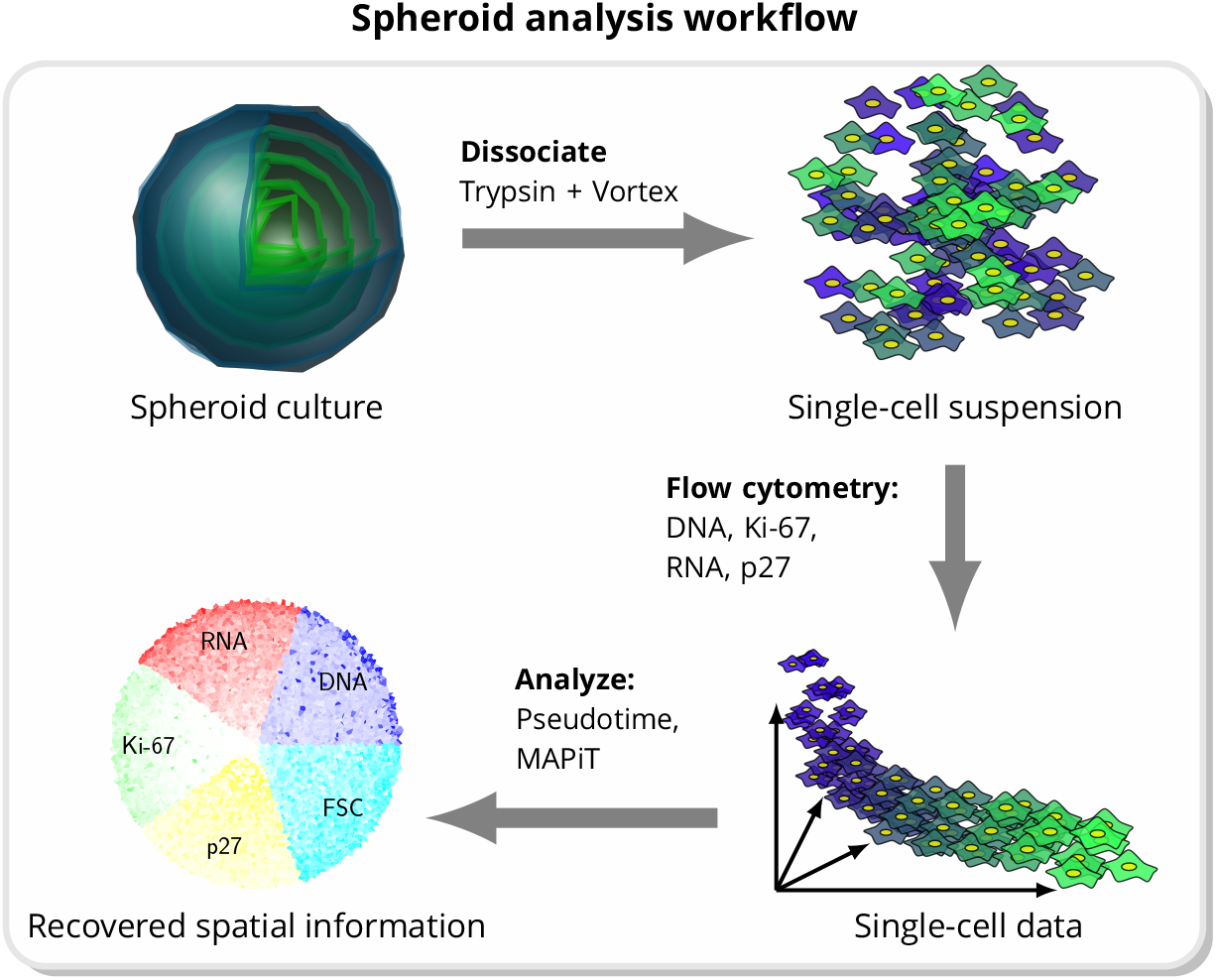
MAPiT recovers spatial positions of cells within spheroids from flow cytometric data. Illustration of spheroid analysis workflow. Individual cells derived from dissociated spheroids were analysed for different markers by flow cytometry. Spatial information can be recovered by applying MAPiT to pseudotime trajectories of measured markers.

To apply MAPiT to these data, the distribution of cells on the spatial scale had to be taken into account, which in the case of radial symmetry of spheroids is the density of cells in relation to the distance from the spheroid surface (Fig. 4 C, Fig. S2). In brief, the volume of a spherical shell on the spheroid surface, normalized to the volume of the whole sphere equals the cumulative distribution of cells in relation to the distance from the surface. A more detailed discussion on spheroid volume and related growth rate is provided in the Supplementary Information.

**Figure 4:**
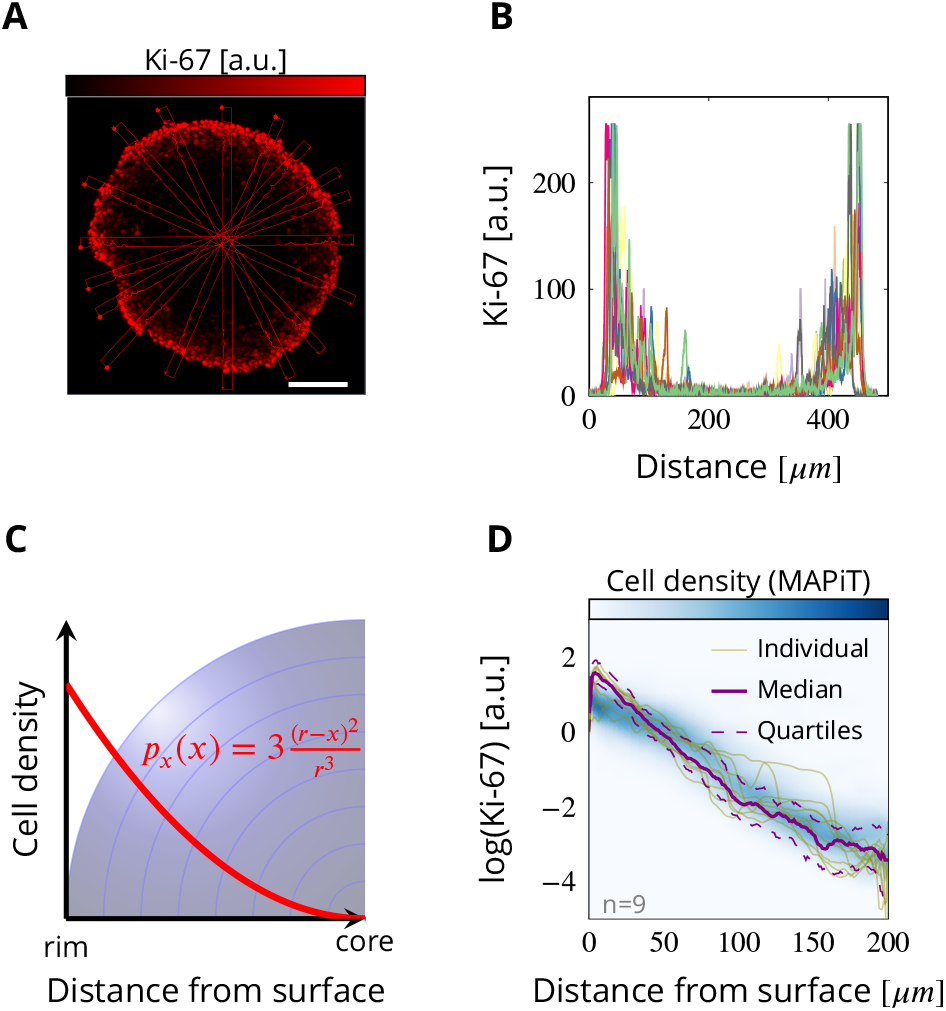
MAPiT derived Ki-67 distribution in an 11-day-old HCT116 cell spheroid. (**A**) Representative spheroid cross-section stained for Ki-67. Scale bar: 100 *μm*. (**B**) Transversal quantified Ki-67 intensities in spheroid sections (rectangles in (i), *n* = 9). (**C**) MAPiT employs cell density in radialsymmetric spheroids with radius *r*. (**D**) Ki-67 intensities obtained by MAPiT from flow cytometric data (blue) closely match Ki-67 intensity profiles determined microscopically in spheroid cross-sections.

MAPiT indeed recovered the spatial positions of single cells, with the reconstructed Ki-67 distribution correlating excellently with intensity profiles obtained from confocal microscopy of spheroid sections stained for Ki-67 (Fig. 4 D). Since MAPiT provides the means to not only employ flow cytometric data but data from any other single-cell based high-throughput multiplex measurement, such as CyToF or single-cell RNA-seq, it provides a foundation for high-throughput and high-content studies of 3D-spheroid models by recovering the spatial information lost during spheroid dissociation (Fig. 3).

Next, we examined if MAPiT can also capture the heterogeneity within the different zones in the spheroid, typically a proliferative layer followed by quiescent and finally necrotic cells.

The DNA content of cells in the outermost spheroid layers exhibited a bimodality typically for proliferating cells with subpopulations in G1 (2N), S and G2/M (4N) phases (Fig. 5 A). In contrast, the majority of cells in the inner layers remain in a quiescent G1/G0 (2N) state. The distinct distributions of additional markers, Ki-67, p27 and RNA content corresponded to this pattern (Fig. 5 A).

**Figure 5:**
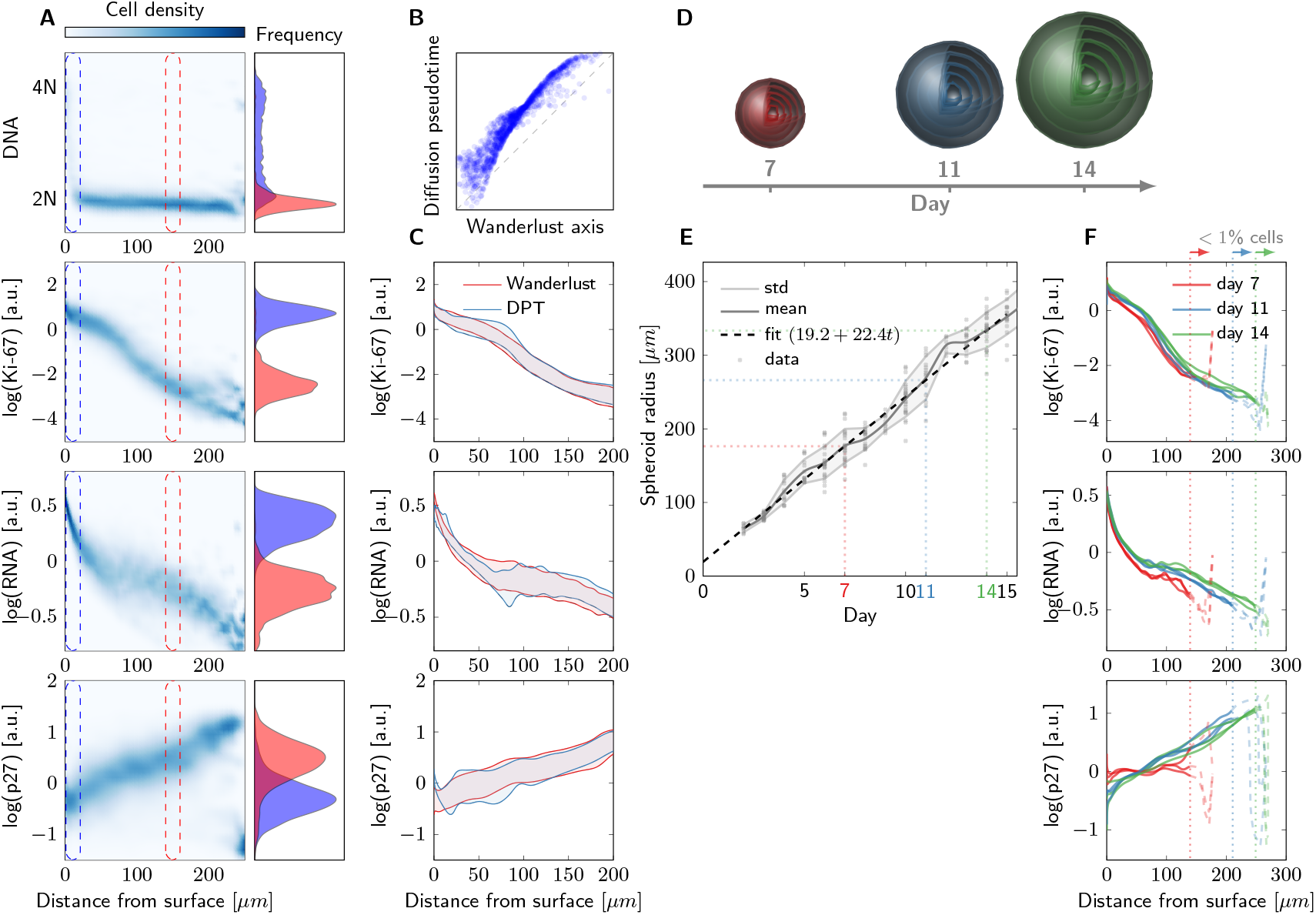
Studying cellular composition in spheroids of HCT116 cells with MAPiT. (**A**) Marginal signal densities *p*(*y|x*) of indicated markers related to the distance from the surface, as obtained by MAPiT. Signal frequencies at the outermost layer and at 150 *μm* distance from the spheroid surface, as indicated by the dashed rectangles, are shown for an exemplary 11-day-old spheroid. (**B**) Comparison of single-cell positions in pseudotime, as obtained by Wanderlust or DPT algorithms. (**C**) MAPiT robustly reconstructs cell positions, irrespective of the pseudotime algorithm used. Ranges show 50%confidence intervals of the signal intensities obtained from transforming Wanderlust and DPT pseudotime data. (**D**) Schematic of spheroid growth. (**E**) Spheroid growth follows a linear growth model. Microscopically obtained data shows spheroid radius over 15 days after seeding from *n* = 23 spheroids in three independent replicates. (**F**) Median profiles for Ki-67, RNA and p27, as obtained by MAPiT, were conserved throughout spheroid sizes (*n* = 3). Notable deviations towards the ends of the profiles (dashed) and thus within the center of the spheroids arise from inaccuracies due to the low number of cells available for analysis at these locations.

We then studied the robustness of MAPiT, by comparing its performance to recover the spatial position using different pseudotime algorithms, markers and spheroid sizes. We first used the Wanderlust and Diffusion Maps algorithms (Bendall et al., 2014; Haghverdi, Buettner, et al., 2014) to obtain a pseudotemporal ordering. The order of cells was largely conserved in both algorithms (Fig. 5 B, monotonously increasing data points), however the pseudotime values were clearly different between both algorithms (Fig. 5 B, deviation from the diagonal). Transforming the DPT and Wanderlust axis to a distance scale with MAPiT resulted in almost identical spatial profiles of Ki-67, RNA and p27 (Fig. 5 C), thereby proving inde-pendence of MAPiT to the pseudotime algorithm. MAPiT should likewise provide identical marker trajectories independent of the choice of markers used to generate the pseudotime order. To validate this, we conducted “leave one out cross validation”, where we took subsets of the markers as input to the Wanderlust and DPT algorithms, and compared the results obtained by MAPiT. MAPiT robustly provided correct distance profiles of all markers in all combinations, demonstrating its independence of the choice of inputs to the pseudotime algorithms (Fig. S1). Thus, MAPiT is independent of the choice of markers or pseudotime algorithms, provided that the order of cells on the pseudotime scale reflects the sequence or directionality of the biological processes.

Spatial reconstruction by MAPiT also provides scope to significantly accelerate high throughput studies that make use of advanced 3D culture-based cell screens, such as spheroid based viability assays and drug effect screens. Indeed, spheroids are considered superior models in comparison to conventional cell cultures (LaBarbera et al., 2012), but spatiotemporal analyses still require cumbersome and manual labor-intensive work for the analysis of crosssectional slices. MAPiT performed reliably in 3D reconstruction when processing flow cytometry data from dissociated spheroids of different sizes and ages, based on Ki-67, RNA and p27 amounts (Fig. 5 D-F). The profiles for Ki-67, RNA and p27 were conserved throughout spheroid sizes, indicating a dependence of these markers solely on the distance from the surface (Fig. 5 E) and therefore their suitability for spatial reconstruction. Overall, robust spatial markers together with MAPiT therefore allow the rapid and versatile reconstruction of spheroids, providing a basis for studying spatiotemporal changes in other measurable variables.

## Discussion

In summary, MAPiT provides a solution to a fundamental problem, namely the transformation of pseudotime to real-time or the true scale of a biological process. For example, snapshot single-cell data of cell populations can be converted to extract real-time kinetics of cellular processes and responses, which otherwise could only be obtained by live-cell microscopy, which is more complex, time consuming and limited by the availability of suitable live-cell reporters. By reconstructing spatial and temporal spheroid compositions from singlecell data, MAPiT provides insights to the evolution of cellular heterogeneity within tumor-like microenvironments and allows to understand how responsiveness to therapeutics manifests within spheroidal environments. By extension, applying MAPiT to other single-cell snapshot data, such as single-cell transcriptomics and proteomics data, might significantly improve the inference of complex regulatory processes and networks by recovering real temporal and spatial dynamics. Recently, pseudotime algorithms were further developed to robustly recognize also branching processes in differentiation pathways (Haghverdi, Büttner, et al., 2016; Setty et al., 2016; Qiu et al., 2017), providing scope to apply MAPiT to study differentiation dynamics in individual branches. Overall, MAPiT is a robust and universal tool to recover temporal or spatial cellular trajectories from high-throughput, high-dimensional single-cell experiments. MAPiT can be combined with pseudotime algorithms, and a MATLAB implementation is available through GitHub (https://github.com/karstenkuritz/MAPiT).

## Materials & Methods

### Calculation of pseudotime

Cells were ordered in pseudotime in MATLAB R2017b (MathWorks, Natick, MA, USA) using two different algorithms: Wanderlust (Bendall et al., 2014) and DPT (Haghverdi, Buettner, et al., 2014). The algorithms were run with 10000 randomly choosen cells. Prior to performing pseudotime analysis, doublets, dead cells and outliers were gated out. Both algorithms require a user defined set of root cells for constructing the pseudotime trajectories. For the cell cycle analysis a set of cells in the center of the G1 population, with low DAPI and geminin signal, was chosen. For the spheroid analysis cells with high Ki-67 and high RNA signal which are known to be located at the surface of the spheroids were chosen.

### MAPiT theory

MAPiT is based on the measure-preserving transformation which states that one must conserve the area under the curve when transforming a probability distribution to another scale.

Consider a measure space 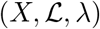, where *X* is a set, 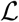 is a *σ* – ring of measurable subsets of *X*, and *λ* is the measure. Given a map *τ* from a measure space 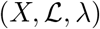 to a measure space 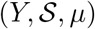, *τ* is called measurable if *A* ∈ *S* implies *τ*^-1^(*A*) ∈ *L*. Given that *τ* is measurable, *τ* is called measure-preserving if *A* ∈ *S* implies *λ*(*τ*^-1^(*A*)) = *μ*(*A*). We denote the pseudotime values with *s* ∈ [0, 1], real-time scale with *x* ∈ [0, *T*] and measured signals with 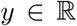. Based on the general definition of a measure-preserving map *τ*, the transformation *τ*: *s* → *x* of a distribution of cells in pseudotime *p_s_*(*s*) to the distribution of cells on the real-time scale *p_x_*(*x*) reads

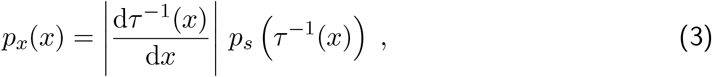

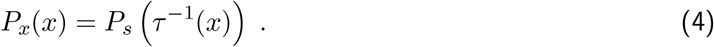

The mapping *τ*: *s* → *x* from pseudotime to real-time was obtained by solving equation (4) for *τ*, which then depends on the cumulative distributions of cells on both scales

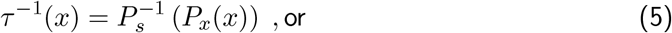

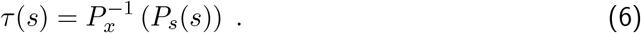

Thus, by definition, the transformation *τ* requires knowledge of the distribution (or cumulative distribution) of cells on the desired scale. Once the mapping *τ* is known, one can apply the transformation to the joint densities of pseudotime and the observed quantities *p_s_*(*s, y*) to obtain the desired joint distribution of the true scale *x* and measured markers *y*

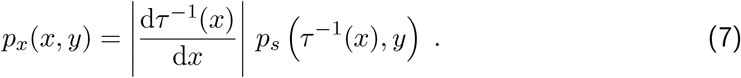

Distributions in pseudotime for spheroid data were obtained by kernel density estimation on pseudotime values *s_i_*, using a Gaussian kernel with reflecting boundary at *s* = {0, 1},

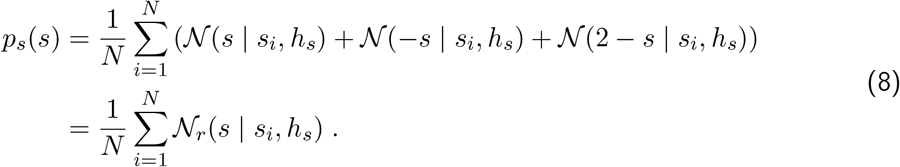

Density in pseudotime for cell cycle data was estimated by kernel density estimation with linked boundary conditions to account for doubling of cell density during cell division (Colbrook et al., 2018). Joint densities were calculated as sum of the product of the individual kernels

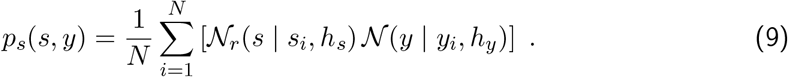

Bandwidths *h_s_* and *h_y_* were derived from Silvermans rule (Silverman, 1986).

### Distributions on the real scales

#### Cell density with respect to cell age in proliferating populations

For analysis of cell cycle-dependent processes with single-cell measurements, MAPiT requires the distribution of cells related to their cell cycle stage or equivalently their age *a*.Cell age, refers to the time since cell birth via cytokinesis. We restricted our analysis to unperturbed cell populations in their exponential growth phase with growth rate *γ* and cell cycle length *T* related by 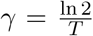. In such a case, the steady state age distribution of a cell population is given by (Powell, 1956)

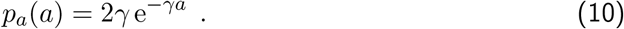

The cumulative distribution and its inverse can be obtained in closed form:

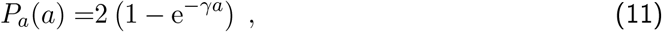

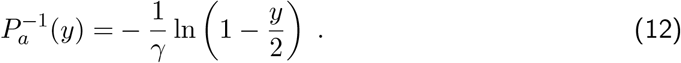

Thus, in case of an unperturbed cell population it is sufficient to know the growth rate of the population to recover cellular age with MAPiT and thus obtain the temporal changes related to cell cycle progression of measured markers from one single-cell experiment.

#### Cell density in tumor cell spheroids with respect to distance from surface

Cell density depending on the distance from the surface was obtained from sphere geometry, assuming radial symmetry (Fig. 2 B, Supplementary Information Fig. S2). The volume of a sphere with radius *r* is given by

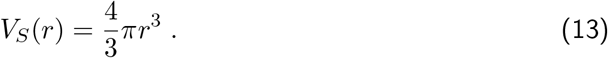

The volume of a spheroid with necrotic core with radius *r_N_* = max(*r* − *d_N_*, 0) equals

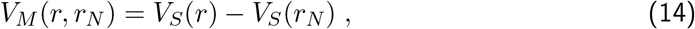

with *d_N_* beeing the distance from the surface where the necrotic core begins. The volume of a spherical shell at distance *x* from the surface of the spheroid with necrotic core is then given by

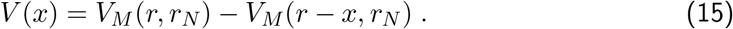

Normalizing (15) with the total spheroid volume *V_M_* results in the normalized volume with respect to the distance to the surface of the spheroid which represents the cumulative distribution of cells related to the distance from the surface

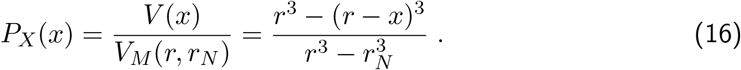

MAPiT is furthermore based on the probability density function and the inverse of the cumulative distribution which can be calculated analytically

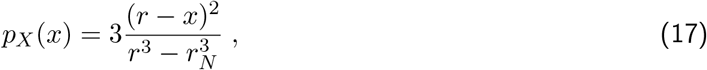

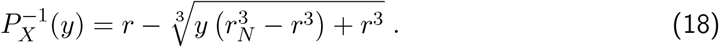

Spheroid radius *r* and the radius of first appearence of a necrotic core *r_N_* were inferred from experiments, described in more detail in supplementary informations. For the present study we used *r_N_* = 270 *μm* and a spheroid radius based on the linear regression

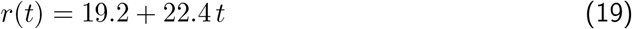

for spheroid growth as shown in Fig. 5 E.

### Cell culture

The human colon carcinoma cell line HCT116 was obtained from the Banca Biologica e

Cell Factory of the IRCCS Azienda Ospedaliera Universitaria San Martino in Genoa (ICLCHTL95025). The geminin expressing non-small-cell lung cancer NCI-H460 cells have been described previously (Kuritz et al., 2017). NCI-H460/geminin cells were maintained in RPMI 1640 medium (Gibco, 21875034) supplemented with 5% heat-inactivated fetal calf serum (FCS, Pan - Biotech GmbH; P303309) and HCT116 cells were cultured in RPMI 1640 medium with 10% heat-inactivated FCS at 37°C in a humidified incubator with 5% CO_2_. For generation of tumor cell spheroids, cells were transferred into Terasaki multiwell plates (Greiner bio-one; 653180) in a volume of 25 *μl* RPMI 1640 medium with a concentration of 4000 cells/ml. Thereafter, the plates were inverted to allow spheroid formation at the bottom of the emerging hanging drops and placed in humid chambers located in the incubator. Two to three days after seeding, formed spheroids were transferred to agarose-coated 96-well cell culture plates (Greiner bio-one; 655180 coated with 1.5% agarose (Carl Roth GmbH & Co. KG; 3810.3) in RPMI 1640 medium). Spheroid growth was monitored with an EVOS FL Cell Imaging System (Thermo Fisher Scientific Inc.) and spheroid diameters were determined from generated pictures using the Fiji software (distribution of ImageJ; Schindelin et al., 2012).

### Time-lapse imaging

NCI-H460/geminin cells were imaged for their total cell cycle length and length of G1 or S/G2/M phases by time-lapse fluorescence microscopy using the Cell Observer system (Carl Zeiss, Oberkochen, Germany) equipped with a humidified imaging chamber at 37°C and 5% CO_2_. Randomly chosen cells were manually tracked for at least one full cell cycle and geminin signal intensity was recorded. Cell trajectories were obtained using the Tracking Tool (tTt) and qTfy for single-cell tracking and quantification of cellular and molecular properties in time-lapse imaging data (Hilsenbeck et al., 2016). For comparison to MAPiT derived results, background signal *θ*_1_ of cell trajectories and scaling factor *θ*_2_ for singlecell trajectories were choosen to maximize the log-likelihood between MAPiT density and individual traces,

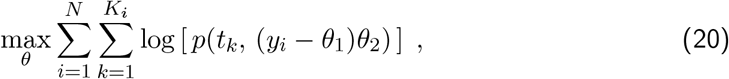

where *p* is the density obtained from MAPiT and *y_i_* are live cell imaging values for geminin of cell *i* at time-points *t_k_* after cell division.

### Generation of spheroid sections and immunofluorescence staining for Ki-67

To generate spheroid sections, spheroids were fixed with 4% paraformaldehyd for 10 min at room temperature. Therafter, spheroids were washed three times with PBS and finally kept in a sucrose solution (30% sucrose (Carl Roth GmbH & Co. KG; 4661) in PBS) at 4°C. After 48 h, the sucrose solution was removed and replaced with Tissue Freezing Medium (A. Hartenstein GmbH; TTEK). Such embedded spheroids were stored at −20°C and finally cut into 10 *μm* slices with a CM30505 cryostat. Generated sections were mounted on Polysine Microscope Adhesion Slides (Thermo Fisher Scientific Inc.; 10219280), fixed with 4% PFA for 10 min at room temperature (RT) and washed twice with PBS for 5 min. Thereafter, permeabilisation was carried out with 0.1% Triton X-100 in PBS for 10 min before the slices were incubated with blocking solution (5% FCS and 0.1% Triton X-100 in PBS) for 30 min at RT. Incubation with the primary antibody against Ki-67 (1:400, Cell Signalling Technology; #9449) was carried out in blocking solution for 1 h at RT. Thereafter, sections were washed two times with blocking solution before they were incubated with the secondary antibody Alexa Fluor 647-conjugated goat anti-mouse IgG (1:500, Thermo Fisher Scientific Inc.; A-21236) solved in blocking solution for 1 h at RT in the dark. Next, sections were washed with blocking solution and incubated with DAPI (1 *μ*g/ml in PBS, Thermo Fisher Scientific Inc.; D1306) to stain DNA for 10 min at RT before they were covered with coverslips using Fluoromount-G^®^ (SouthernBiotech; 0100-01). Samples were dried and fluorescence was analysed with a LSM 710 laser scanning microscope (Carl Zeiss, Oberkochen, Germany) and the blue edition of the ZEN software.

### Flow cytometric analysis

Tumor cell spheroids were dissociated with trypsin/EDTA (Gibco; 59418C) and single cells were fixed with 4% PFA in PBS for 10 min. Thereafter, permeabilisation was carried out with 90% ice-cold methanol in PBS for 30 min on ice. Subsequently, cells were washed two times with a washing solution (5% FCS, 0.05% BSA (Sigma-Aldrich; A2058), 0.02% NaN_3_(Carl Roth GmbH & Co. KG; Ca4221) in PBS) before they were incubated with the primary antibodies diluted in washing solution for 1 h at RT (anti-Ki-67, Cell Signalling Technology; #9449 and anti-p27, Cell Signalling Technology; #3686). Next, cells were washed and incubated with the respective secondary antibodies, Alexa Fluor 488-conjugated goat anti-rabbit IgG and Alexa Fluor 647-conjugated goat anti-mouse IgG diluted in washing solution for 1 h (1:100, Thermo Fisher Scientific Inc.; A11008 and 1:500, Thermo Fisher Scientific Inc.; A21236). After washing, cells were incubated with 100 *μl* Hoechst 33342 solution (10 *μg/ml* Hoechst 33342 (Thermo Fisher Scientific Inc.; H3570) in PBS with 0.05% BSA and 0.02% NaN_3_) for 1 h at 37°C, followed by the addition of 5 *μl* Pyronin Y (stock solution: 100 *μg/ml* solved in ddH_2_O, Sigma-Aldrich; 83200) and incubation for 15 min at 37^°^C. Finally, cells were pelleted, dissolved in PBS and fluorescence was measured by flow cytometry with the MACSQuant analyser 10. NCI-H460/geminin cells were analyzed as described in Kuritz et al., 2017.

## Acknowledgments

We would like to thank the German Research Foundation (DFG) for financial support of the project within the Cluster of Excellence in Simulation Technology (EXC 310/2) at the University of Stuttgart.

## Author contributions

K.K. designed and implemented the mathematical framework. K.K., D.S., D.M. and N.P. designed and carried out the experiments. K.K., D.S, and N.P. analyzed and interpreted the data. M.R. and F.A. supervised the project; K.K., D.S., N.P., M.R., and F.A. wrote the paper with feedback from all authors.

