## Supplementary Information for "Reconstructing temporal and spatial dynamics in single-cell experiments"

Karsten Kuritz[1,*], Daniela Stöhr[2], Daniela Maichl[2], Nadine Pollak[1,2,3], Markus Rehm[2,3,+] and Frank Allgöwer[1,3,+]

[1]Institute for Systems Theory and Automatic Control, University of Stuttgart, Stuttgart, Germany

[2]Institute of Cell Biology and Immunology, University of Stuttgart, Stuttgart, Germany

[3]Stuttgart Research Center Systems Biology, University of Stuttgart, Stuttgart, Germany

[+]shared senior author

### Supplementary Material

#### Cross-validation of marker selection

We examined how MAPiT is affected by the choice of observations used to generate the pseudotime order. Therefore, MAPiT results were compared in a "leave one out cross validation" type approach, where we took subsets of the observations as input to the pseudotime algorithm. Of the four measured markers (DNA, Ki-67, RNA and p27) subsets of only two markers were used to obtain a pseudotime ordering with pseudotime algorithms like Wanderlust and DPT. Only Ki-67, RNA and p27 are relevant for cell ordering, resulting in three distinct combinations: RNA+Ki-67, RNA+p27 and Ki-67+p27. Despite small deviations at the beginning and the end of the pseudotime, both algorithms found the same order of cells with all marker combinations (Supplementary Fig. S1 A). A conserved order consequently entails high similarity in the profiles verifying the independence of MAPiT to the choice of observations (Supplementary Fig. S1 B).

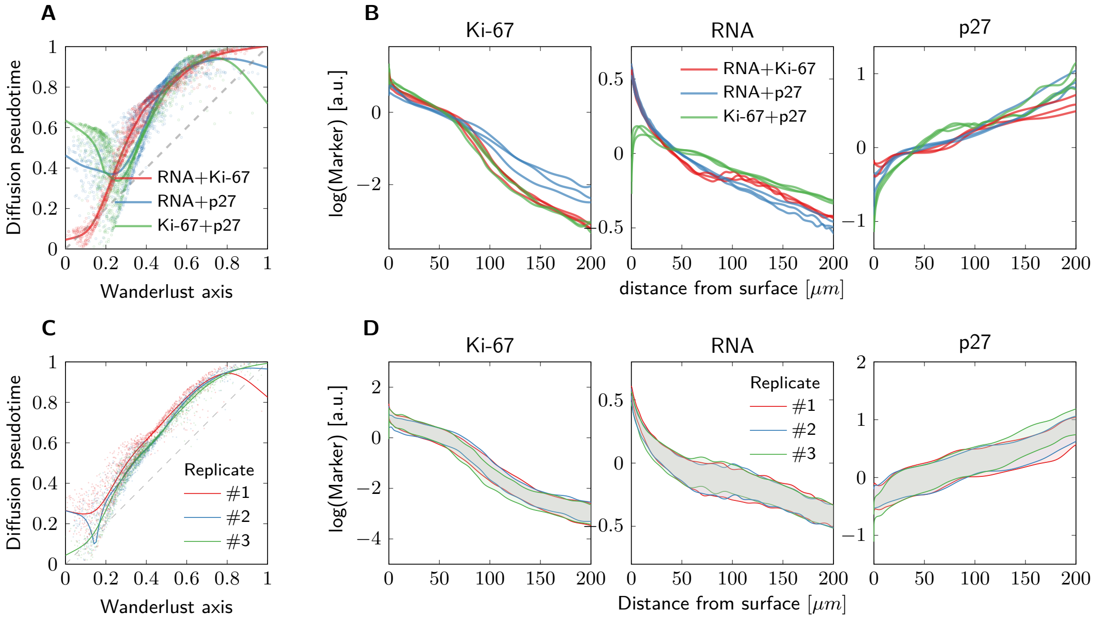


**Figure S1:** Robust reconstruction of distance-dependent signal intensities from observation subsets in 11-day-old HCT116 spheroids. (**A**) Single-cell Wanderlust axis position vs. Diffusion pseudotime position for subsets of the four measure markers. (**B**) Three replicates of median intensities of the transformed signal obtained by performing Wanderlust with indicated markers. (**C**) Wanderlust axis position vs. DPT position of spheroid-derived samples from three biological replicates. (**D**) $50\%$ confidence intervals of the signals transformed from pseudotime, obtained by Wanderlust, of three replicates.

### Spheroid growth dynamics

In order to correctly calculate the transformation from pseudotime to the distance scale the cumulative density of cells in the spheroid w.r.t. the distance from the surface must be known. This requires (1) the size of the spheroid and (2) the size at which the necrotic core emerges (Fig. S2). Both quantities can be obtained from models of spheroid growth dynamics.

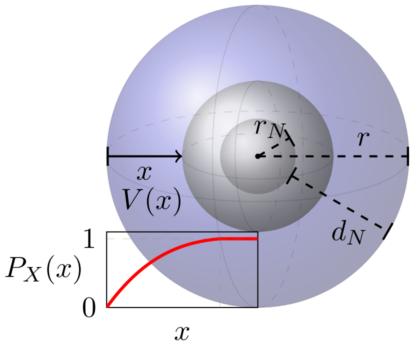


**Figure S2:** Sphere geometry. Volume of spherical shell $V(x)$ with radius $r$ and thickness $x$ (blue), normalized to the volume of the sphere $V_S(r)$ minus the volume of the necrotic core $V_S(r - r_N)$ provides the cumulative distribution of cells in the spheroid $P_X(x)$.

The spheroid volume as a function of time can be calculated with a spheroid growth model with solely one layer of proliferative cells at the spheroid surface (Conger and Ziskin, 1983). However, emergence of a necrotic core is not captured with such a simple model but has to be estimated from experimental data. Spheroids are seen as analogous to avascular tissue or tumor mass and diffusion limitation to many molecules, particularly $O_2$, becomes apparent in spheroids with diameters larger $150 - 200\,\mu m$ (Lin and Chang, 2008). In addition, inefficient mass transport leads to metabolic waste accumulation inside the spheroids. Therefore, spheroids with a diameter larger $500\,\mu m$ commonly display a layer-like structure comprising a necrotic core surrounded by a viable rim. The viable rim consists of an inner layer of quiescent cells and an outer layer of proliferating cells (Lin and Chang, 2008; LaBarbera et al., 2012). We developed a layer-based growth model to identify the spheroid size at which the necrotic core emerges. The model consists of $i \in \{1, \ldots, I\}$ different layers of which each layer has its own characteristics determined by the parameters of the layer:

- $d_i$ layer thickness in $\mu m$
- $\gamma_i$ fraction of proliferating cells in layer $i$
- $\mu_i$ growth rate in $\frac{1}{\text{day}}$ of the proliferating cells in layer $i$

Furthermore we have two parameters that are independent of the layers:

- $r_0$ initial spheroid radius in $\mu m$.
- $v_c$ volume of a single cell in $\mu m^3$.

We defined the innermost layer as a necrotic core without any cells. The necrotic core then emerges at a spheroid radius which is equal to the sum of the estimated thickness of the outer layers

$$r_N = \sum_{i=1}^{I-1} d_i \ . \tag{1}$$

Parameters $d_i, \gamma_i, \mu_i$ thus are sufficient to describe the growth of a spheroid. The volume growth of a spheroid is given as the sum of the volume change in each layer by:

$$\begin{aligned} \frac{\mathrm{d}V}{\mathrm{d}t} &= \sum_{i=1}^{I} V_i \gamma_i \mu_i \\ &= \frac{4}{3}\pi \sum_{i=1}^{I} \gamma_i \mu_i \left( (r - d_{i-1})^3 - (r - d_{i-1} - d_i)^3 \right) \ . \end{aligned} \tag{2}$$

The change of the the volume can furthermore be decomposed with the chain rule into

$$\frac{\mathrm{d}V}{\mathrm{d}t} = \frac{\mathrm{d}V}{\mathrm{d}r}\frac{\mathrm{d}r}{\mathrm{d}t} \ . \tag{3}$$

We know that the change of the volume with respect to the radius is given by

$$\frac{\mathrm{d}V}{\mathrm{d}r} = 4\pi r^2 \ . \tag{4}$$

By inserting (2) and (4) into (3) one solves for a change of the radius of the spheroid over time, which is sufficient to describe the whole spheroid growth

$$\frac{\mathrm{d}r}{\mathrm{d}t} = \frac{1}{3r^2} \sum_{i=1}^{I} \gamma_i \mu_i \left( (r - d_{i-1})^3 - (r - d_{i-1} - d_i)^3 \right) \ . \tag{5}$$

This nonlinear ODE (5) can be solved numerically to obtain the spheroid radius over time $r(t)$. Other quantities of interest can then be computed:

- Spheroid volume: $V(t) = \frac{4}{3}\pi r(t)^3$
- Total cell number in the spheroid: $N(t) = \frac{V(t)}{v_c}$
- Volume of the single layers: $V_i(t) = \frac{4}{3}\pi \left( \left( r(t) - \sum_{j=1}^{i-1} d_j \right)^3 - \left( r(t) - \sum_{j=1}^{i} d_j \right)^3 \right)$
- Cell number per layer: $N_i(t) = \frac{V_i(t)}{v_c}$
- Number of proliferating cells in a layer: $N_i^P(t) = N_i(t)\gamma_i$
- Number of quiescent cells in a layer: $N_i^Q(t) = N_i(t)(1 - \gamma_i)$

#### Parameter estimation for the spheroid growth model

The parameters of the spheroid growth model were identified by comparing the model predictions to spheroid volumes obtained by microscopy and cell numbers obtained from flow cytometry counts of individual spheroids at different days post seeding. We used a weighted sum of squares as a measure for the difference between the dataset and the model predictions. The parameters were estimated initially using the CMA-ES algorithm (Hansen, 2006). In a next step the parameters were sampled using a MCMC DRAM algorithm to additionally obtain the uncertainties of the parameter values (Haario et al., 2006). The models were initiated with different numbers of layers ranging from $I \in \{2, 3, 4\}$. Based on the Bayesian Information Criterion (BIC), the model with two layers followed by a necrotic core ($I = 3$) was identified as the most plausible. The spheriods then consisted of an outer proliferating layer with thickness of $20 - 25\,\mu m$, which is close to the estimated diameter of a single cell. The second layer has a thickness of roughly $250\,\mu m$ and does not contain proliferating cells. The necrotic core emerges at a radius of $r_N \approx 270$ which is in good agreement with literature values (Lin and Chang, 2008; LaBarbera et al., 2012). Based on the estimated parameters a reduced ODE model for the spheroid radius is given by:

$$\frac{\mathrm{d}r}{\mathrm{d}t} = \mu_1 \left( d_1 - \frac{d_1^2}{r} - \frac{d_1^3}{r^2} \right) . \tag{6}$$

Once the spheroid reaches a certain size where $r >> d_1$ Eq.(6) reduces to a constant growth of the spheroid

$$\frac{\mathrm{d}r}{\mathrm{d}t} = \mu_1 d_1 \ , \tag{7}$$

which reflects the simple linear growth model of the spheroid radius already presented in (Conger and Ziskin, 1983). Our layer-based model thus could reproduce experimental data and is furthermore in line with previous spheroid growth models. The derived values for spheroid growth $r(t)$ and the radius at which the necrotic core emerges $r_N$ were used to calculate the position-dependent cell density,

$$p_X(x) = 3\frac{(r-x)^2}{r^3 - r_N^3} \ , \tag{8}$$

required for the application of MAPiT to single-cell data of dissociated spheroids.

#### Comparison to other methods

To allow for a profound discussion of MAPiT and other methods, we first describe the underlying mathematical description of 1D processes related to the transformation of pseudotime to real-time. The evolution of the distribution of cells on the process manifold $s \in [0, 1]$ can be described by a second order partial differential equation

$$\frac{\partial}{\partial t}n(s,t) + \frac{\partial}{\partial s}\left(v(s)n(s,t)\right) - \frac{1}{2}\frac{\partial^2}{\partial s^2}\left(g(s)^2 n(s,t)\right) = \left(\alpha(s) - \beta(s)\right) n(s,t) \ , \tag{9}$$

with position-dependent velocity and diffusion $v(s)$ and $g(s)$, respectively. Source $\alpha(s)$ and sink $\beta(s)$ capture growth of the population, e.g., by cell division and vanishing cells, e.g., due to cell death. Neumann boundary conditions describe influx and outflow at the beginning

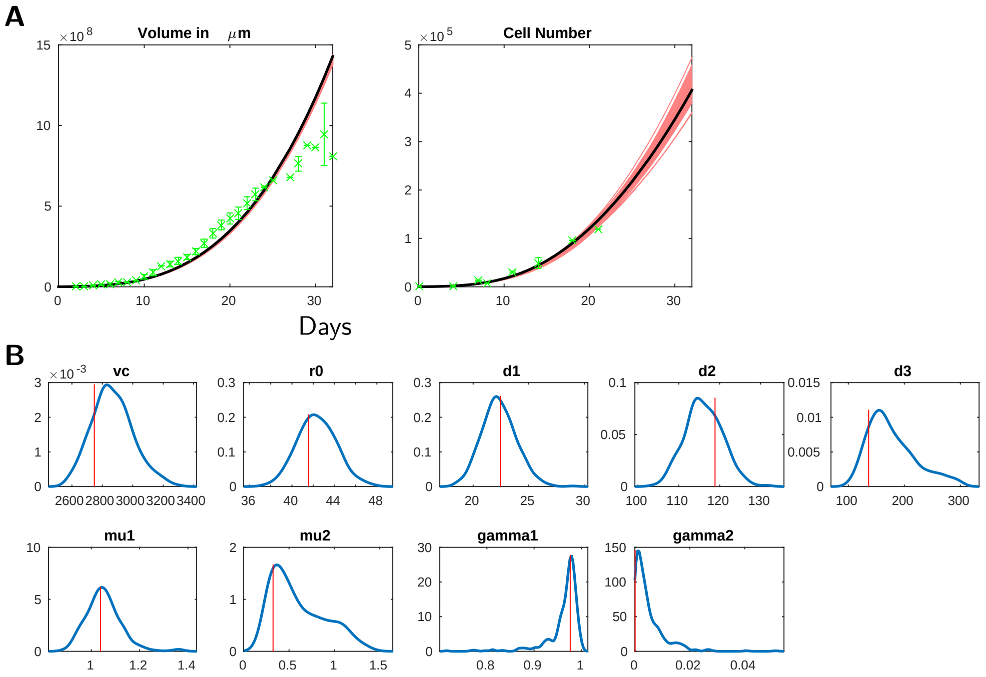


**Figure S3:** Parameter estimation results for HCT116 spheroid growth kinetics. (**A**) Predictions of a three layer model with a necrotic core for spheroid volume and total cell number. Best fit (black) and 200 best samples (red) in comparison with experimental data (green). Model predictions correspond very well with the measurements. (**B**) Posterior parameter distributions and best fit (red) obtained by MCMC of a three layer spheroid growth model. Parameter *vc* and *r0* represent single cell volume and spheroid radius at $t = 0$. Parameters d1-d3 define thickness of layers with distinct properties such as growth rates *mu1*, *mu2* and fraction of proliferating cells *gamma1*, *gamma2* in the corresponding layer. All parameters were identifiable.

and the end of the process

$$\begin{aligned}\frac{\partial}{\partial s}n(0,t) &= J_0(t)\\ \frac{\partial}{\partial s}n(1,t) &= J_1(t)\ ,\end{aligned} \tag{10}$$

and an initial condition for the distribution at time $t = 0$ completes the model

$$n(s,0) = n_0(s)\ . \tag{11}$$

Few methods allow the transition from pseudotime to real-time analysis: *Pseudodynamics* infers real-time dynamics by estimating the distribution of a cell population across a continuous cell state coordinate over time based on a stochastic differential equation (Fischer et al., 2019). Estimated functions are not uniquely identifiable and the method is computationaly demanding which might impair its applicability. Another approach called *velocyto* is based on the abundance of different RNA splicing variants restricting the method to scRNA-Seq data (La Manno et al., 2018). Moreover, ergodic theory on cyclic processes was used to infer the rates of molecular events during cell cycle progression from single-cell measurements (Kafri et al., 2013; Kuritz et al., 2017).

Haghverdi and colleagues suggested inference of a universal time by integrating over the inverse distribution in pseudotime (Haghverdi et al., 2016). The theoretical basis behind this is that the density on the pseudotime scale is inversely correlated with the velocity, with which the cells progress through this position on the process manifold. Given the PDE eqs. (9) to (11), this holds true if and only if

$$\alpha(s) - \beta(s) = 0\ , \tag{12}$$

$$g(s) = 0\ , \tag{13}$$

$$J_0(t) = J_1(t)\ . \tag{14}$$

This is equal to assuming that, (1) population growth (by cell division) and cell removal (by cell death) balance each other, (2) all cells are identical and (3) influx of cells in the beginning of the process is equal to the outflow at the end. While (2) and (3) are under certain conditions reasonable assumptions, balancing of population growth and cell death (3) at each stage of the process is only justified in special cases. However, if (1)-(3) hold and the process is in steady state operation, then one can deduce, based on ergodic principles, that the derived universal time is the same as the real-time.

In another approach by Weinreb and colleagues, Population Balance Analysis (PBA) was used to infer diffusion $g(s)$ and entry and exit rates $\alpha(s), \beta(s)$ in a PDE similare to eqs. (9) to (11) (Weinreb et al., 2018). However, the recovered time scale was still not identical to real-time due to nonidentifiability of velocity and combined entry and exit rates at the same time.

Pseudodynamics takes a similar approach to fit functions of the PDE (9) to the observed distributions from snapshot data (Fischer et al., 2019). By using a regularization Fischer and colleagues try to overcome the nonidentifiability of velocity and combined entry and exit rate.

The method called velocyto uses a different approach (La Manno et al., 2018). Local velocity is inferred from the abundance of splicing variants at each point in the data space. By assuming a conserved model with transcript independent mRNA splicing rates, one obtains velocity information directly from the abundance of different splicing variants in scRNA-Seq data. The concept is very elegant, however the method is restricted to single-cell transcriptomics and the assumption of constant rates might not always hold.
